# Uveal melanoma cells use ameboid and mesenchymal mechanisms of cell motility crossing the endothelium

**DOI:** 10.1101/2020.03.18.997627

**Authors:** Michael D. Onken, John A. Cooper

## Abstract

Uveal melanomas (UM) are malignant cancers arising from the pigmented layers of the eye. UM cells spread through the bloodstream, and circulating UM cells are detectable in patients before metastases appear. Extravasation of UM cells, notably transendothelial migration (TEM), is a key step in formation of metastases. UM cells execute TEM via a stepwise process of intercalation into the endothelial monolayer involving the actin-based processes of ameboid blebbing and mesenchymal lamellipodial protrusion. UM cancers are driven by oncogenic mutations in Gαq/11, which activate TRIO, a guanine nucleotide exchange factor (GEF) for RhoA and Rac1. Pharmacologic inhibition of Gαq/11 in UM cells reduced TEM. Inhibition of the RhoA pathway blocked amoeboid motility but led to enhanced TEM; in contrast, inhibition of the Rac1 pathway decreased mesenchymal motility and reduced TEM. Inhibition of Arp2/3 also inhibited mesenchymal motility, but invasion was less affected; in this case, the amoeboid blebbing behavior of the cells led to transmigration without intercalation, a direct mechanism similar to that of immune cells. BAP1-deficient (+/−) UM subclones displayed motility behavior and increased levels of TEM, similar to effects of RhoA inhibitors. We conclude that RhoA and Rac1 signaling pathways, downstream of oncogenic Gαq/11, combine with pathways regulated by BAP1 to control the motility and transmigration of UM cells.

## INTRODUCTION

Uveal melanoma (UM) is a highly aggressive cancer arising in the uveal layers of eye, including the choroid, ciliary bodies, and iris. The eye is the second most common site of melanoma. UM and cutaneous melanoma are distinct diseases [1, 2]. Oncogenesis of UM tumors is driven by constitutive activation of the Gq family of alpha subunits, Gαq and Gα11 (*GNAQ* and *GNA11*) in over 90% of tumors, while cutaneous melanomas are driven by *BRAF* and *NRAS* mutations [3, 4]. Ten-year survival of patients with primary UM tumors is about 50%, compared to over 90% for cutaneous melanoma [5]. The anatomy of the eye essentially prevents local spread, and the posterior chamber of the eye lacks lymphatic vessels, so metastatic spread of UM to distant organs occurs only through the bloodstream [6]. Despite local control of the primary eye tumor being achieved in over 95% of patients, UM has a high rate of metastasis and lethal outcome [7].

Hematogenous spread of UM cells begins with shedding of cells from the primary eye tumor. The tumor vasculature of UM is irregular and discontinuous, providing a poor barrier to cell shedding [8, 9]. As a result, circulating tumor cells are found in all patients, even those with low-risk tumors that do not develop metastatic disease [10–12]. Because shedding of cells into the blood is common, extravasation of circulating tumor cells out of blood vessels at distant sites may be a key rate limiting step for UM metastasis.

To investigate transendothelial migration (TEM), we developed a cell culture system employing primary human endothelial monolayers grown on polyacrylamide hydrogels, and we followed TEM in real time with living cells [13]. Using this approach, we discovered that the migration of UM cells occurs via a stepwise process [14]. Suspended UM cells attach to endothelial cells and then intercalate between adjacent endothelial cells, flattening into the monolayer. UM cells remain intercalated for up to several hours before releasing from the endothelial monolayer and migrating underneath it. The cellular and molecular mechanisms driving each step have not been characterized and may provide information for preventing circulating tumor cells from exiting the vasculature.

Cell migration and the actin cytoskeleton are key features of UM metastasis. Spread of UM tumors to distant organs is driven by loss of BAP1, a chromatin remodeling factor [15], and we found that depletion of BAP1 increases the overall rate of TEM in our system [14]. BAP1 is a histone deubiquitinase recruited to promoters of genes [16], and we found that several actin cytoskeleton regulator genes are targeted by BAP1 in UM cells [16]. In parallel, constitutively active Gq/11, the oncogenic driver of UM, activates the dual nucleotide exchange factor TRIO, which in turn activates the Rho-like small GTPases RhoA and Rac1 [17] that regulate the actin cytoskeleton during cell migration [18, 19]. We hypothesize that regulation of actin-based processes, downstream of BAP1 and Gq/11, and via the RhoA and Rac1 pathways, controls the migration of UM cells.

Rac1 drives actin polymerization at the leading edge of a migrating cell and promotes focal complex assembly [20, 21], resulting in the formation of lamellae and lamellipodia, which move the cell forward [20–22]. Rac1 activates downstream effectors, including WAVE2, which activate Arp2/3 complex for branched actin polymerization [23]. Rac1 also activates p21-activated kinase (PAK), which has several important substrates [24, 25], including GEF-H1, a regulator of RhoA [26].

RhoA activates Rho-associated protein kinases 1 and 2 (ROCK1 and ROCK2) [27, 28], which increase actomyosin contractility via myosin light-chain (MLC) phosphorylation. This results in increased cortical contractile forces, which enhance membrane blebbing [29]. Biochemically, the RhoA and Rac1 pathways can be antagonistic, with opposing roles in cell migration [26, 28]. ROCK signaling can antagonize Rac1 via ARHGAP22, and Rac1-WAVE2 activity can antagonize ROCK signaling [28]. Interacting and antagonistic pathways downstream of RhoA and Rac1 have been identified in normal fibroblasts and breast cancer [30, 31].

Here, we investigated the roles of these signaling pathways with a combination of pharmacologic and molecular genetic perturbations, using direct live-cell observation of TEM. We found that UM cells in suspension show predominantly amoeboid motility driven by the RhoA pathway, and that UM cells switch from amoeboid motility to mesenchymal motility as they adhere to and migrate through the endothelial monolayer.

## RESULTS

First, we established methods to quantify UM cell behavior during TEM, allowing us to measure the effects of pharmacologic and genetic perturbations. We characterized the movements of 92.1 UM cells before and during TEM from live-cell movies. UM cells displayed amoeboid blebbing while in suspension, immediately after addition to endothelial monolayers, and for a short while after coming to rest on the monolayers. The cells continued blebbing during their initial interactions with the endothelial monolayer. Next, they stopped blebbing and formed lamellipodial protrusions. The cells proceeded to spread, flatten, and intercalate themselves between endothelial cells (Figure 1A, and Video S1). The steps of adhesion and intercalation occurred relatively quickly and were complete within a few hours. After a longer period of time, the cells formed new lamellipodial protrusions that extended underneath the endothelial monolayer, and the cells migrated away, in agreement with our previous observations [14].

**Figure 1.**
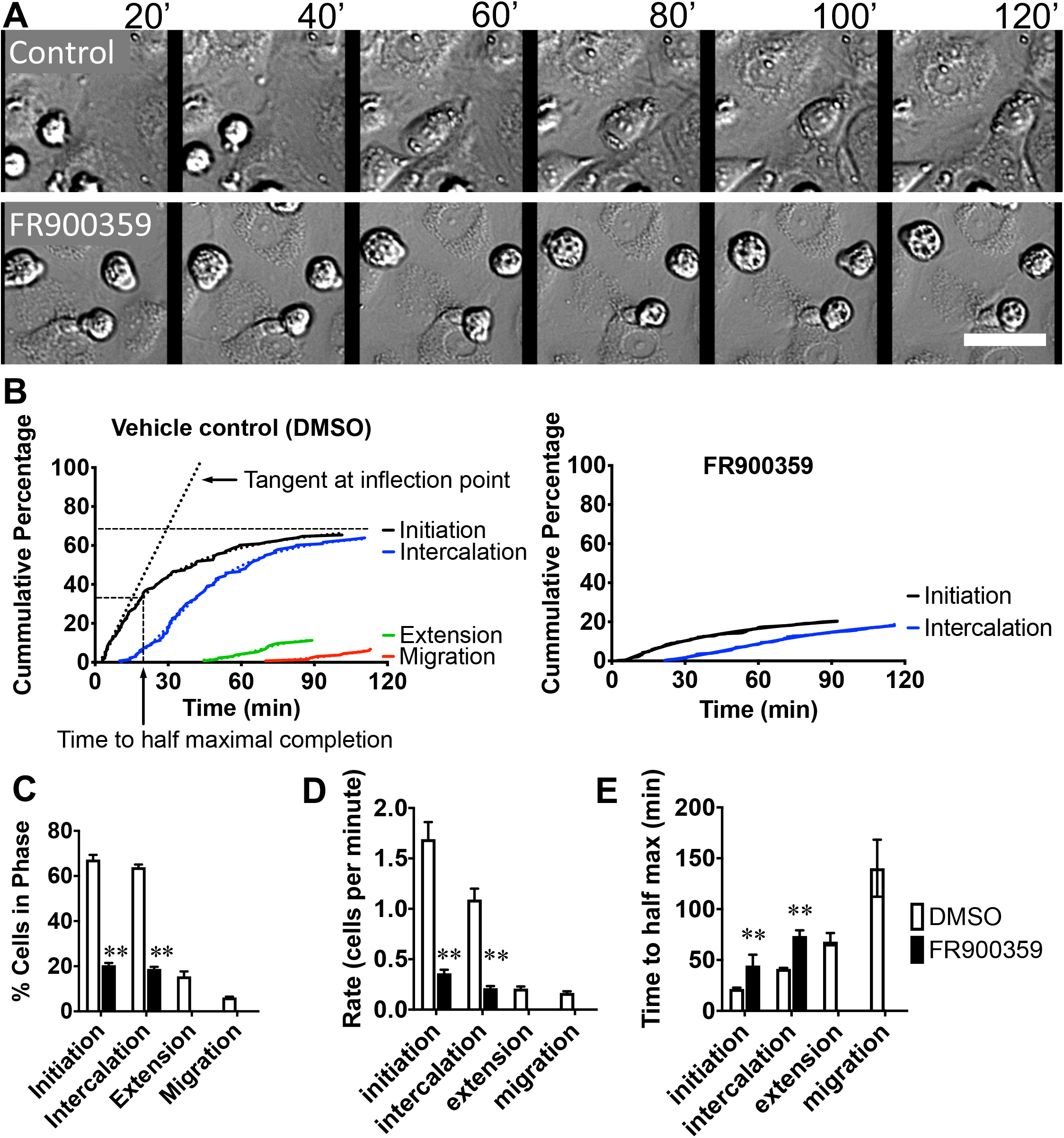
Quantification of TEM of untreated and FR900359-treated UM cells. **A.** Frames from a representative movie of TEM by control UM cells (upper panels, Video S1) and cells treated with FR900359 (lower panels, Video S2), at 20-min intervals. Scale bar = 50 *μ*m. **B.** Cumulative percentage of cells undergoing each step of UM cell transmigration. Results are from three 2-hr movies for each condition. Each curve was quantified to generate the bar graphs in panels C-E (p < 0.05, *; p < 0.01, **). **C.** Endpoint values for each step after 2 hours. **D.** Maximum rate of each step calculated from the slope of the tangent at the inflection point. **E.** Time to reach half of the calculated maximum plateau for each step.

We quantified the progression of UM cells through each of these steps during TEM (see Figure 1B for an example) by measuring and calculating the following parameters: a) total number of cells completing each step after 2 hours (Figure 1C), b) maximum instantaneous rate of cells completing each step (Figure 1D), and c) time required for half the expected number of cells to complete each step (Figure 1E). The endothelial monolayers were prepared from primary HDMVECs, for which different lots display a certain level of biological variability. To account for this variability among lots, control (DMSO-treated) samples were performed with every experiment, and the results from these control samples were used to normalize the results from experimental samples.

### Inhibition of the Oncogenic Driver blocks TEM

Constitutively activating mutations in Gq/11 are the driving oncogenes for over 90% of UM tumors [32]. Gq/11 activates the dual-GEF TRIO, which activates RhoA and Rac1 in UM cells [18]. We used FR900359, a potent inhibitor of Gq/11 in UM cells [33] to block TRIO activation by Gq. UM cells treated with FR900359 for 24 hrs showed significant decreases in progression for all steps of transmigration (Figure 1A and Video S2). Adhesion and intercalation were reduced to 40% of untreated activity, and neither extension nor migration were detected during any 2-hour experiment (Figure 1C). Maximum rates of progression of each step decreased by >4-fold (Figure 1D), time to reach maxima increased by >2-fold (Figure 1E). Thus, inhibition of the oncogenic driver mutation blocks most UM cells from transmigrating and slows the progression of the few cells that do transmigrate.

### Inhibition of the RhoA pathway increases TEM

Activated TRIO regulates both RhoA and Rac1, which control opposing cell morphology pathways [28]. We interrogated each pathway independently to dissect their regulatory roles in UM cells during TEM. Inhibition of ROCK, the immediate downstream effector of activated RhoA, by Y-27632 blocked the amoeboid blebbing activity of UM cells in suspension and on the surface of the monolayers (Figure 2A and Video S3). Surprisingly, UM cells treated with Y-27632 showed significant increases in both the percentage of cells completing each step of TEM (Figure 2B-C) and the rates of cells progressing through each step (Figures 2D-E).

**Figure 2.**
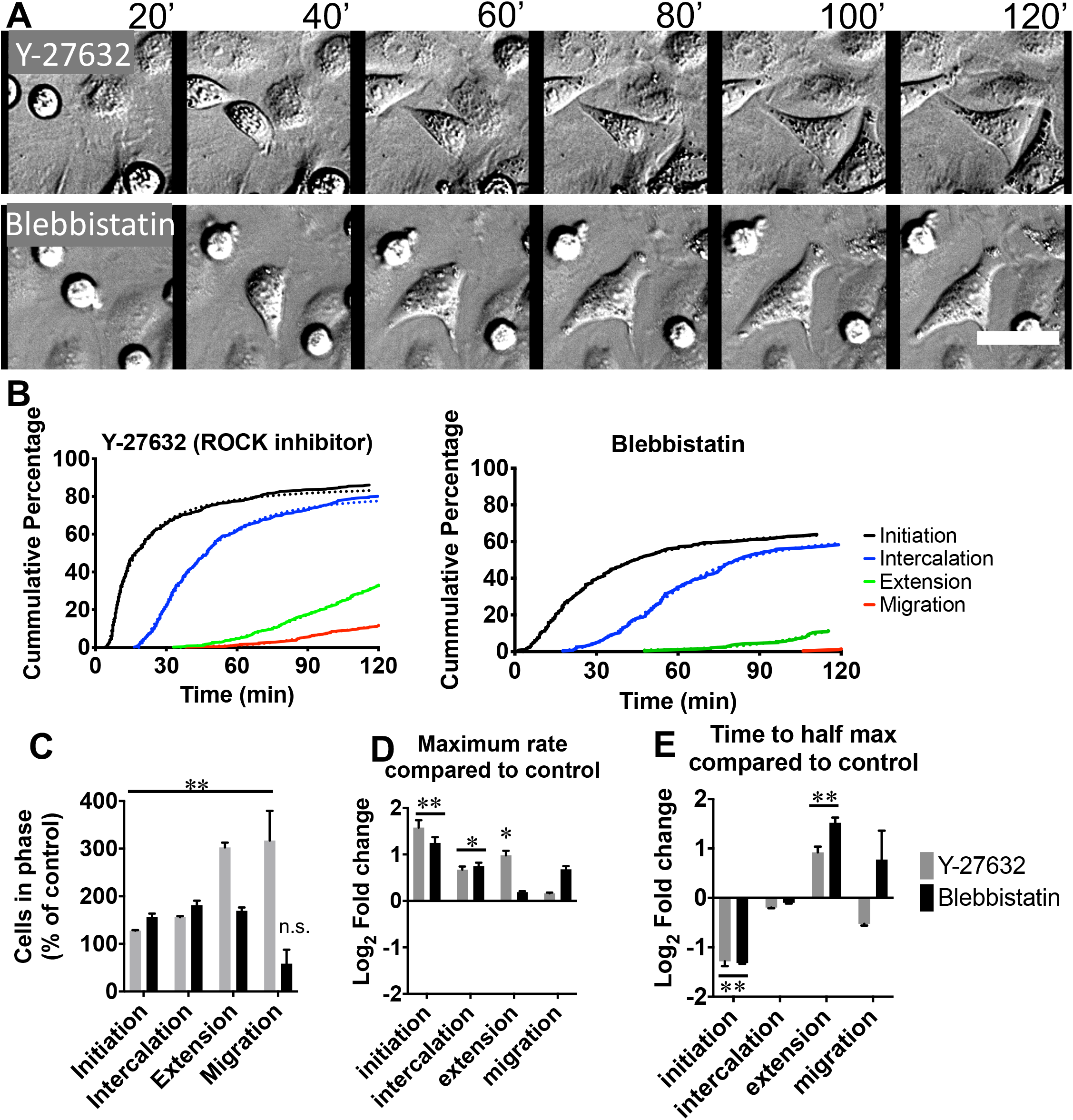
Effects of ROCK and myosin inhibition on TEM of UM cells. **A.** Inhibition of RhoA-dependent ROCK1 by Y-27632 (upper panels, Video S3) or myosin by blebbistatin (lower panels, Video S4). **B.** Cumulative percentage of cells that have undergone each step of UM cell transmigration, based on results from three 2-hr movies. **C.** Normalized endpoint values of data in panel B. **D.** Normalized maximum rates for each step of TEM. **E.** Normalized times to reach half of the calculated maximum plateaus. Treatment with either inhibitor causes more UM cells to transmigrate and at a faster rate. Scale bar, 50 *μ*m; p < 0.05, *; p < 0.01, **.

Downstream of ROCK in the RhoA pathway is non-muscle myosin II, the motor protein that powers cellular contractility and membrane tension and thus drives amoeboid blebbing [34]. Inhibition of non-muscle myosin II by blebbistatin blocked blebbing activity in UM cells (Figure 2A and Video S4) and significantly increased the rates of TEM of UM cells (Figure 2B-E). These effects were similar to those of Y-27632. Thus, the effects of ROCK inhibition on UM cells appear to occur through regulation of actomyosin contractility.

### Inhibition of the Rac1 pathway blocks TEM

Actin-based lamellipodial protrusions are a prominent feature of UM cells as they intercalate into the endothelial monolayer and then extend out of the monolayer and migration underneath it. We hypothesized that these protrusions were driven by activation of Rac1. To inhibit Rac1, we treated UM cells with NSC-23766, which blocks the activation of Rac1 by exchange factors. The cells showed increased amoeboid blebbing and decreased lamellipodial protrusion, with an overall impairment of TEM (Figure 3A-B and Video S5). The numbers of cells completing each step were significantly reduced, as were the rates of completion within each step (Figure 3C-E). The decreases in the TEM parameters were similar to the strong effects on cells treated with FR900359 above (Figure 1C-E).

**Figure 3.**
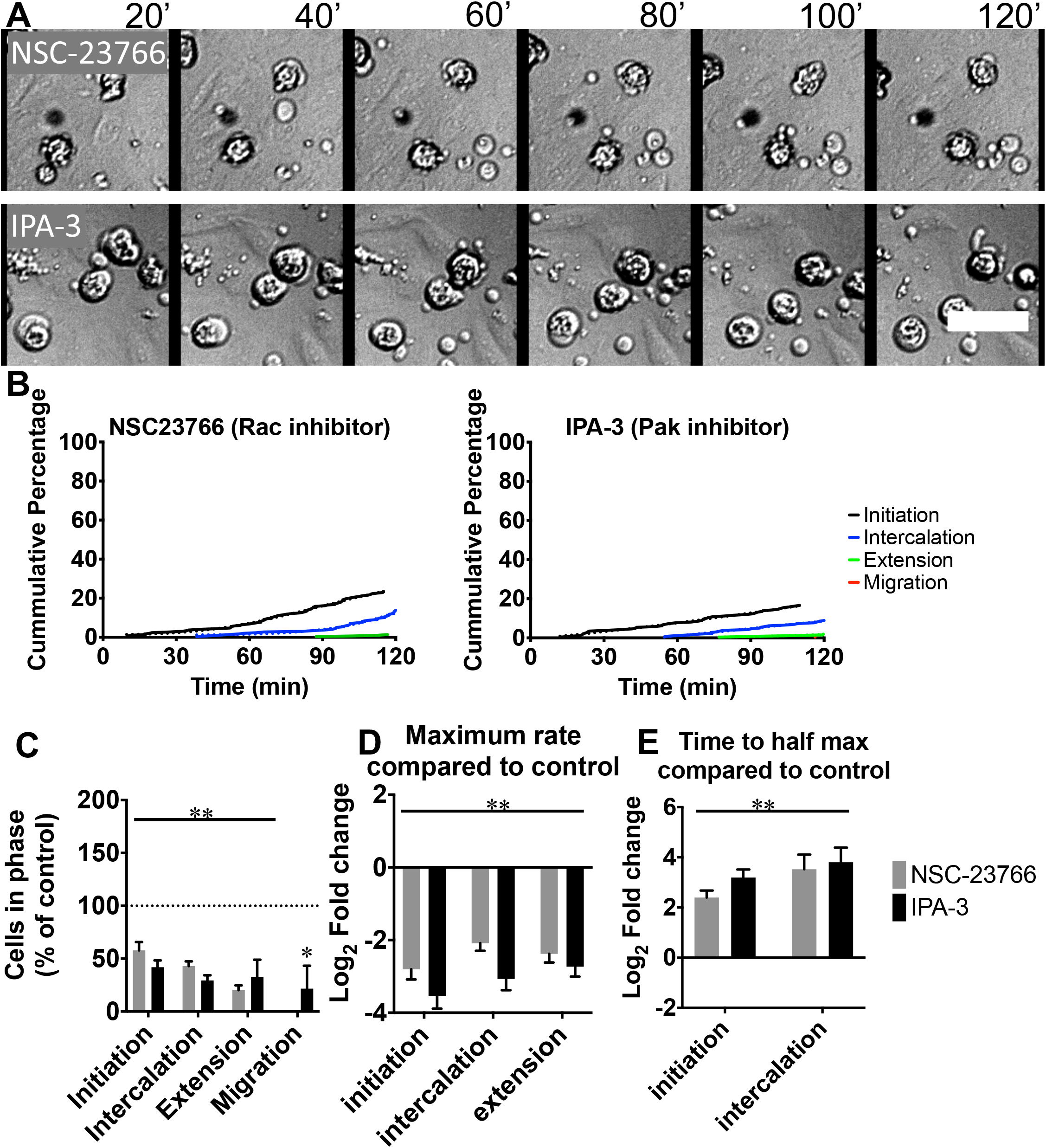
Effects of Rac1 and PAK inhibition on TEM of UM cells. **A.** The RacGEF inhibitor NSC-23766 (upper panels, Video S5) and the PAK inhibitor IPA-3 (lower panels, Video S6) inhibit all steps of TEM. **B.** Cumulative percentage of cells that have undergone each step based on results from three 2-hr movies. **C.** Normalized endpoint values of panel B. No cells treated with NSC-23766 migrated underneath the monolayer after 2 hours. **D.** Normalized maximum rates for each step. The number of TEM events was too low to calculate maximum rates of migration. **E.** Normalized times to reach half of the calculated maximum plateaus. Treatment with either inhibitor caused fewer UM cells to transmigrate and at a slower rate. The number of TEM events was too low to accurately analyze extension and migration. Scale bar, 50 *μ*m; p < 0.05, *; p < 0.01, **.

Rac1 directly activates PAKs, which phosphorylate target proteins [25]. Treatment of UM cells with the PAK inhibitor IPA-3 enhanced blebbing and reduced TEM (Figure 3A-B), similar to the effects of NSC-23766. Fewer cells completed each step, and rates of completion within each step were reduced (Figure 3C-E). Interestingly, UM cells treated with either NSC-23766 or IPA-3 also showed larger blebs and increased extracellular debris (Figure 3A and Videos S5 and S6) compared to untreated cells (Figure 1A and Video S1), suggesting dysregulation of bleb size and retraction resulting from inhibition of the Rac pathway.

### Inhibition of Arp2/3: UM cells transmigrate and bypass intercalation

The actin-based protrusions that occur during lamellar and lamellipodial cell migration are driven by the rapid formation and growth of branched actin networks at the cell periphery. Actin branching is nucleated by Arp2/3 complex, which is activated by the WAVE complex, a downstream target of activated Rac1. We used the Arp2/3 inhibitor CK-666 [35] to interrogate the role of branched actin formation in Rac1 regulation of TEM. UM cells treated with CK-666 showed extensive blebbing, similar to the effects of NSC-23766 and IPA-3 (Figure 4A and Video S7). Despite failing to switch from amoeboid blebbing to mesenchymal motility, CK-666-treated UM cells proceeded to perform TEM in a direct manner, bypassing the process of intercalation (Figure 4A and Video S8). This TEM migration behavior is similar to that of immune cells, including lymphocytes and NK cells [13].

**Figure 4.**
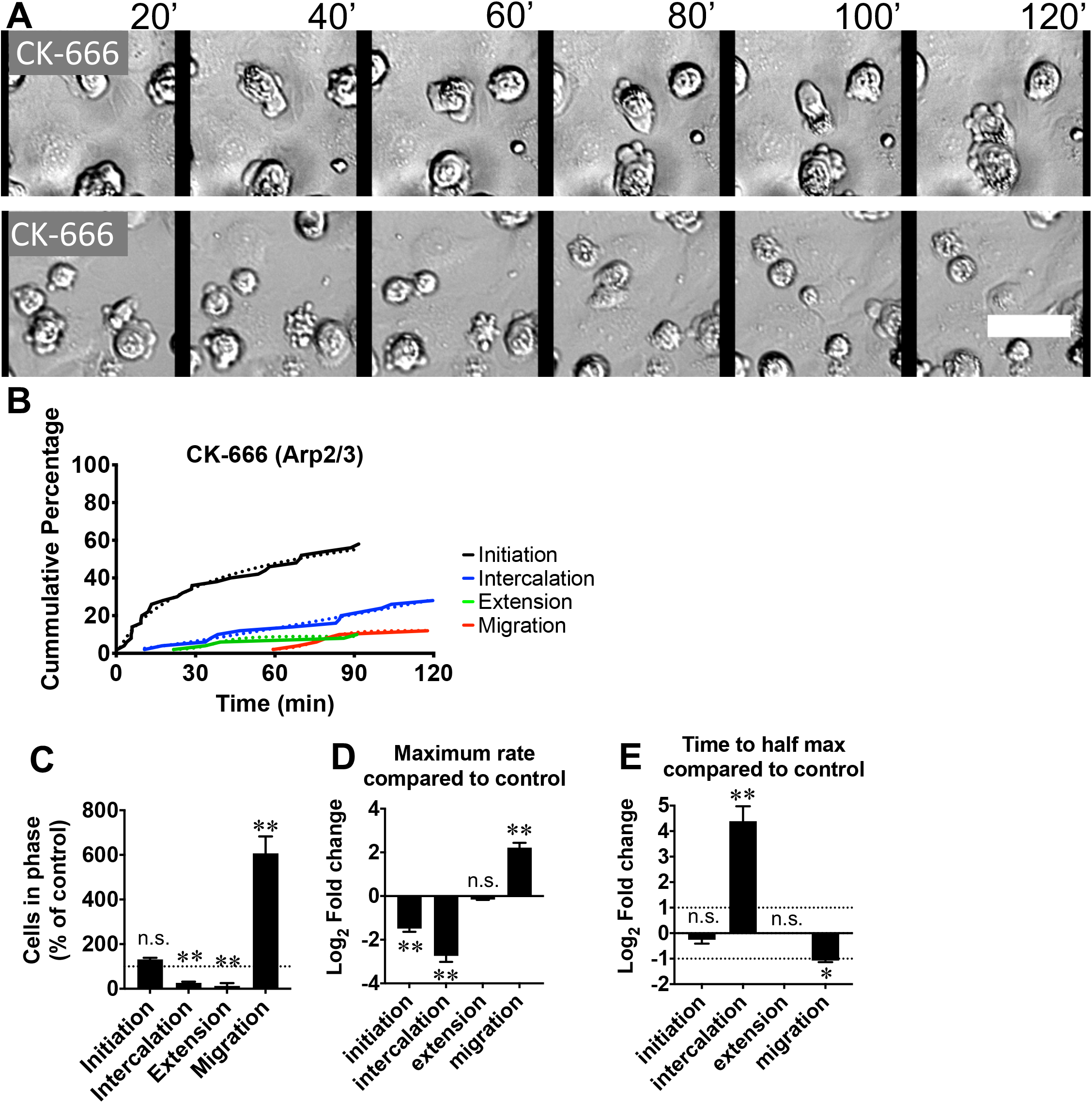
Effects of Arp2/3 inhibition on TEM of UM cells. **A.** CK-666 blocked the switch to lamellipodial activity, decreasing intercalation (upper panels, Video S7) and increasing direct transmigration bypassing intercalation (lower panels, Video S8). **B.** Cumulative percentage of cells that have undergone each step of transmigration. **C.** Normalized endpoint values of panel B. **D.** Normalized maximum rates were decreased for intercalation but increased for migration. **E.** Normalized times to reach half of the calculated maximum plateaus. Scale bar, 50 *μ*m; p < 0.05, *; p < 0.01, **.

### Depletion of the Metastasis Suppressor BAP1 increases TEM

Loss of the chromatin-remodeling factor BAP1 drives metastasis in UM tumors [15, 36]. BAP1 regulates the transcription of TRIO, cortactin, GTPase activating proteins (GAPs) and guanine nucleotide exchange factors (GEFs) in UM (Figure S1) [16], which could alter the balance of RhoA and Rac1 pathways. We used CRISPR/Cas9 genome editing to generate BAP1-deficient UM cells. While the homozygous loss of BAP1 was lethal in all of several UM cell lines tested, we were able to generate two independent BAP1-deficient heterozygous subclones of the 92.1 UM cell line, 92.1-1D7 and 92.1-2D3. Both BAP1(+/−) UM cell lines displayed amoeboid blebbing while in suspension, and they switched to forming lamellipodial protrusions during intercalation, extension, and migration, similar to parental BAP1(+/+) cells (Figure 5A and Videos S9, S10). However, both subclones showed quantitative increases in all steps of TEM, with more cells completing all steps, and with significantly faster times for initiation and intercalation than parental BAP1(+/+) cells (Figure 5C-E). Thus, reduction of BAP1 activity produced overall effects on TEM similar to those resulting from inhibition of the RhoA pathway as opposed to the effects of inhibition of the Rac1 pathway (Figure S2). These data suggest that BAP1 expression favors activation of the RhoA pathway over the Rac1 pathway, such that loss of BAP1 shifts the balance toward Rac1-driven motility and increased TEM.

**Figure 5.**
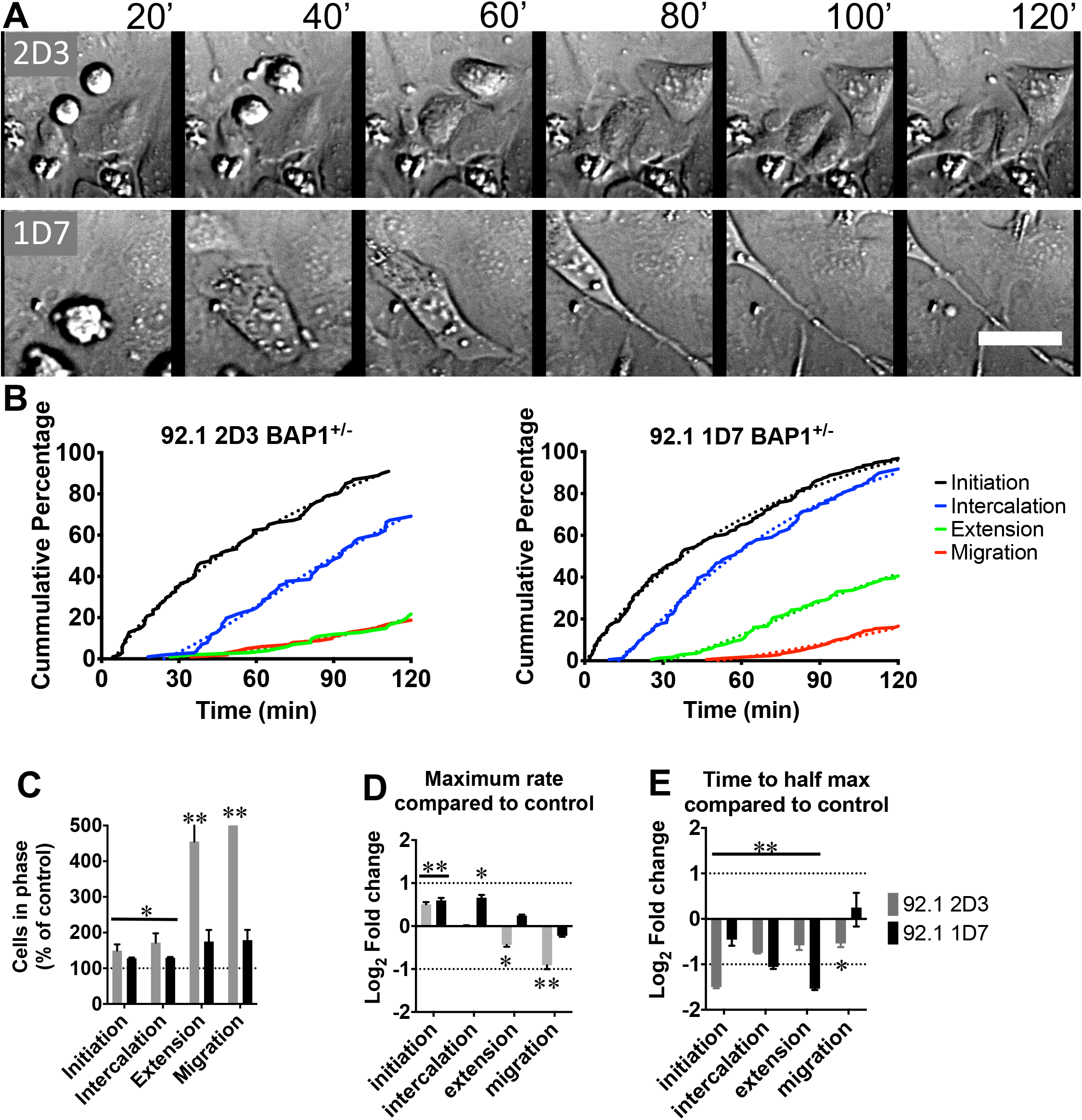
BAP1-deficient UM cells show increases in all steps of TEM. **A.** Frames from time-lapse movie of two independent BAP1(+/−) subclones of the UM 92.1 cell line: 2D3, (upper panels, Video S9) and 1D7 (lower panels, Video S10). **B.** Cumulative percentage of cells that have undergone each step based on results from three 2-hr movies. **C.** Normalized Endpoint values of panel B. **D.** Normalized maximum rates. **E.** Times to reach half of the calculated maximum plateaus. Scale bar, 50 *μ*m; p < 0.05, *; p < 0.01, **.

## DISCUSSION

### UM cells switch modes of motility during TEM

We discovered that UM cells exhibit both amoeboid and mesenchymal modes of motility as they migrate through the endothelium during TEM. UM cells perform TEM by a stereotypical and characteristic multi-step route, which includes intercalation of the UM cell into the endothelial monolayer [14]; here, we discovered that their ability to switch from amoeboid to mesenchymal motility is essential to intercalation. Previous studies observed amoeboid and mesenchymal morphologies for cutaneous melanoma cells attached to substrates with varying levels of adhesivity [28, 37–39]; those studies did not involve melanoma cells interacting with the endothelium.

We discovered that the oncogenic driver mutation of UM cells is responsible for promoting TEM. Inhibition of constitutively active Gq/11 in UM cells, by the pharmacologic inhibitor FR900359, led to a nearly complete loss of TEM activity in our assay system. In our previous work, FR900359 caused UM cells to stop dividing and to re-differentiate towards their melanocytic state [33]. Together, the results suggest that FR900359 has promise as a novel therapeutic agent for UM tumors. The inhibition of TEM observed here may be particularly significant because the lethality of UM tumors is due to metastasis through the bloodstream [6, 10] and thus requires TEM.

We discovered that the Rho and Rac pathways activated downstream of the oncogenic mutation in UM cells have distinct complementary roles in the mechanism of transmigration. Inhibiting the Rho pathway blocked amoeboid motility but increased TEM (Figure S2), while inhibiting the Rac pathway blocked mesenchymal motility and decreased TEM (Figure S2).

We propose the following model of TEM regulation in UM cells (Figure 6). UM cells display amoeboid morphology and motility as they flow through the bloodstream and make initial contact with the endothelium. This amoeboid behavior depends on active RhoA and actomyosin contraction. Following contact with the endothelium, UM cells convert their morphology and motility to Rac1-driven lamellar and lamellipodial protrusions. These protrusive actions enhance the interactions of UM cells with the endothelial cells, allow the UM cells to intercalate and flatten into the endothelium, and then leave the endothelium and migrate into surrounding tissues.

**Figure 6.**
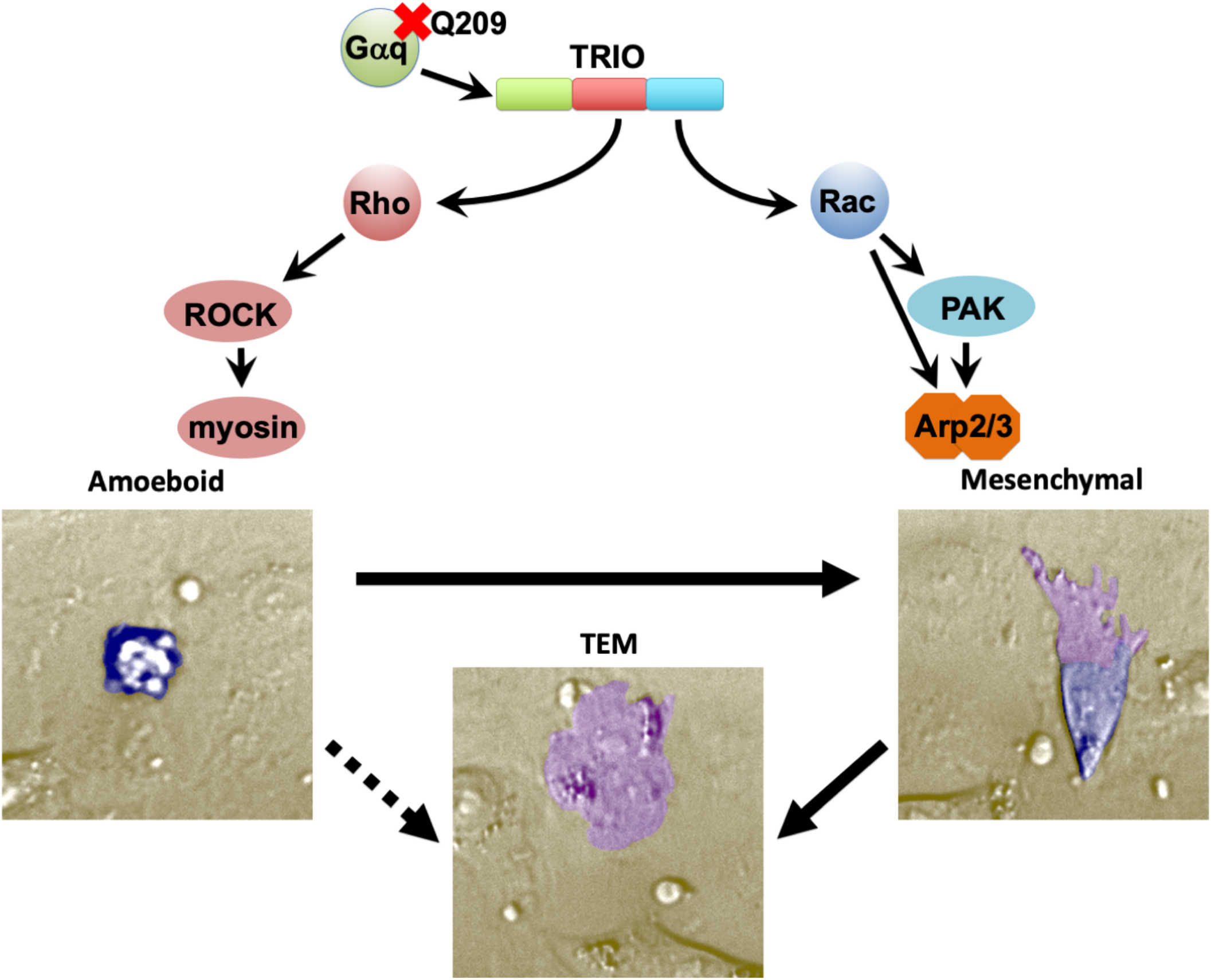
Diagram Summarizing UM Cell Signaling and Modes of Motility during TEM. Constitutively active Gq, the oncogenic driver for UM, signals through the dual-GEF TRIO, which activates both RhoA and Rac1. Rho signals through ROCK to myosin-II, which promotes amoeboid motility. Rac activates PAK and Arp2/3, which promote lamellar and lamellipodial protrusions as part of mesenchymal motility. The switch from amoeboid to mesenchymal motility increases the robustness of TEM, but TEM can occur via a solely amoeboid mechanism.

### Amoeboid and Mesenchymal Migration: Complementary Roles for TEM

UM cells treated with Arp2/3 inhibitor interacted with the endothelial monolayer and performed TEM, but they did not switch from the amoeboid mode to the mesenchymal mode of motility and migration. In this case, the intercalation step was dispensable for TEM, and amoeboid behavior alone was sufficient. This finding suggests that UM cells in situ may use a combination of amoeboid and mesenchymal modes of migration, depending on the local circumstances, such as the architecture of the vascular bed and the flow of the bloodstream. One might speculate that inside-out signaling from Rac and PAK activates adhesion molecules on the UM cell surface, such as integrin α4β1 [40], required for adhesion to endothelial cells [14]. This hypothesis is supported by our observation of decreased interactions of Rac/PAK-inhibited UM cells with the endothelial cell monolayer, compared to untreated or Arp2/3-inhibited UM cells.

Another possible explanation for the effect of Arp2/3 inhibition is a requirement for branched actin assembly to stabilize early interactions that promote the maturation of adhesive junctions. One model of cell-cell adhesion formation is that weak interactions at a small cell-cell contact surface recruit active Rac1, which activates actin assembly to expand the contact surface and form a mature adhesion [41]. This model is based on observations that Arp2/3 inhibition disrupts junction formation between cells on plastic but not the formation of cell aggregates. Cells that aggregate with neighboring cells do not require extensive Arp2/3-dependent lamellipodia activity, but cells that are flat and spread out require expanded contacts through lamellipodia, which would require Arp2/3 activity [42]. This model predicts that blocking branched actin formation by inhibiting Rac1 or PAK would also prevent UM cells from establishing sufficiently strong adhesions with the endothelial monolayer. Our results expand this view by suggesting that separate adhesive mechanisms exist, dependent on and independent of stabilization by actin assembly during TEM.

### Metastasis-promoting BAP1 mutation favors mesenchymal motility and increases transmigration

We discovered that partial loss of BAP1, based on heterozygous gene deletion, caused increases in all steps of TEM. This discovery is important because in human UM patients, the loss of BAP1 is the key genetic feature associated with a high risk of metastasis in the class 2 lethal genotype [15, 36]. The clinical relationship of deletion of BAP1 with metastatic spread is consistent with the biological connection of BAP1 function with the mechanism of TEM discovered here. The genomic transcriptional targets of BAP1 include a number of actin cytoskeleton genes, including TRIO, cortactin, and various GAPs and GEFs (Figure S1A) [16]. BAP1 targets of particular note include *ARHGAP18*, a Rho-specific GAP, and *ARHGEF4*, a Rac/cdc42-specific GEF [16], both of which are upregulated in BAP1-deficient Class 2 human tumors [43] (Figure S1B). Concurrent upregulation of ARHGAP18 and ARHGEF4 would be expected to decrease RhoA signaling and increase Rac1 signaling and promote TEM. In support of this hypothesis, increased expression of ARHGAP18 [44, 45] and ARHGEF4 [46] have been linked to metastasis in other cancers.

## ACKNOWLEDGMENTS

This work was supported by NIH grant GM118171 to J.A.C.

## AUTHOR CONTRIBUTIONS

Conceptualization, M.D.O. and J.A.C.; Methodology, M.D.O. and J.A.C.; Investigation, M.D.O.; Formal Analysis, M.D.O.; Visualization, M.D.O.; Writing - Original Draft, M.D.O.; Writing - Review & Editing, M.D.O. and J.A.C.; Funding Acquisition, J.A.C.; Resources, M.D.O. and J.A.C.; Supervision, M.D.O. and J.A.C.

---

***Figure S1: Transcriptional regulation of Rho and Rac pathways by BAP1.*** Reanalysis of published data sets. **A.** BAP1 genomic targets identified by calling-card results [16]. Black bars indicate number of transposon insertions and white bars indicate significance of peak height at transcription start sites for each gene. **B.** Gene expression in human tumor samples [43]. Class 1 (blue) are characterized by normal BAP1 and low metastasis probability, and Class 2 (red) are characterized by BAP1 gene deficiency and high metastasis probability. Each dot represents an expression value for one human patient tumor sample.

***Figure S2: Summary of Results for RhoA and Rac1 Pathways, revealing opposing effects on TEM.*** Combination of all the data in Figures 1-5, for comparison. **A.** Normalized maximum rates of cells that have undergone each step. **B.** Times to reach half of the calculated maximum plateaus.

## METHODS

### RESOURCE AVAILABILITY

#### Lead Contact

Further information and requests for resources and reagents should be directed to and will be fulfilled by the Lead Contact, Michael Onken.

#### Materials Availability

This study did not generate new unique reagents.

#### Data and Code Availability

This study did not generate any novel datasets.

### EXPERIMENTAL MODEL AND SUBJECT DETAILS

Cells were cultured at 37°C in a humidified 5% CO_2_ incubator. Human 92.1 UM cells (RRID: CVCL_8607) were the generous gift of Dr. Martine Jager (Laboratory of Ophthalmology, Leiden University, Netherlands) and were grown in RPMI 1640 medium (Life Technologies, Carlsbad, CA) supplemented with 10% FBS and antibiotics. Primary human dermal microvascular endothelial cells (HDMVECs: HMVEC-dNeo, Catalog #CC-2516; Lonza, Allendale, NJ) were grown in EGM-2 MV culture medium (Lonza). HDMVECs were not used after 8 passages. BAP1(+/−) 92.1 subclone cell lines were generated using CRISPR/Cas9 genome editing and sequence-verified by the Washington University Genome Engineering and iPSC Center (http://geic.wustl.edu).

### METHOD DETAILS

#### Reagents and Inhibitors

Chemicals and reagents were obtained from Fisher Scientific (Pittsburgh, PA) or Sigma-Aldrich (Saint Louis, MO), except as follows: Blebbistatin, Y-27632, NSC-23766, and IPA-3 were from Selleck Chemicals (Houston, TX). FR900359 was the generous gift of Dr. Kendall J. Blumer (Department of Cell Biology & Physiology, Washington University, St. Louis, MO). Inhibitors were dissolved in DMSO as stock solutions and frozen in small aliquots. Inhibitors were tested for toxicity using a tetrazolium-based Cell Counting Kit-8 (Bimake, Houston, TX) at working concentrations for 12 hrs. The inhibitors did not significantly reduce cell viability, with the exception of FR900359, which has long-term effects that have been described elsewhere [33]. For TEM experiments, Blebbistatin and Y-27632 were used at 10 *μ*M, IPA-3 was used at 30 *μ*M, CK-666 was used at 100 *μ*M, and FR900359 was used at 100 nM.

#### Live Cell Imaging

Polyacrylamide hydrogels were prepared with 0.4% bis-acrylamide on glass-bottom microwell dishes (MatTek corp., Ashland, MA) as described [13]. The hydrogels were coated with fibronectin (10 *μ*g/mL in PBS) by incubation at 37°C for 30 minutes. HDMVECs were plated onto the fibronectin-coated hydrogel substrates and incubated overnight to allow formation of monolayers. Monolayers were inspected by phase-contrast microscopy to ensure that cells covered the substrate completely, without defects, before transmigration assays were performed. Monolayers were rinsed with fresh EGM-2 MV media to remove floating cells and debris one hr before assaying TEM.

UM cells were trypsinized, collected in medium containing FBS to stop tryptic activity, then centrifuged and resuspended in EGM-2 MV culture medium at 10^5^ cells / mL. Inhibitors or vehicle were added to individual aliquots of UM cells two hrs prior to TEM imaging, except for FR900359, which was added to stock plates one day before imaging (>24 hours) and again after harvesting and resuspension in EGM-2 MV. For imaging, HDMVEC monolayers were placed into an environmental chamber (Stage Top Incubator, Tokai Hit, Shizuoka-ken, Japan) with 5% CO_2_ at 37°C on an inverted microscope (Olympus IX72), and 10^4^ UM cells were added to the monolayer directly over the objective. DIC images were captured at 20-s intervals with a 10X objective. To control for biological variability among the primary human endothelial cells, DMSO-treated experiments were performed with every inhibitor experiment, and those results were used for normalization.

### QUANTIFICATION AND STATISTICAL ANALYSIS

Statistical analyses were performed with Prism (GraphPad, San Diego, CA). For endpoint cell counts, mean and standard error of the mean (SEM) were calculated from at least three experiments for each inhibitor, and statistical analysis compared each inhibitor experiment to control experiments from the same day. To generate cumulative percentage curves, data from all experiments for a given inhibitor were combined, and these curves were used to quantify rates. Non-linear regression was used to fit curves for each step, which extrapolated to maximal completion plateau values used to quantify time to half maximal completion. To calculate maximum rates, tangent lines were drawn either at the most vertical points of curves that showed clear change over time or along the longest linear stretch for curves that followed a more linear course. Times and rates for each step were normalized to values generated from control experiments run on the same day and 90% confidence intervals were calculated.

## REFERENCES

1. Urtatiz O., Cook C., Huang J.L., Yeh I., and Van Raamsdonk C.D. (2020). GNAQ^Q209L^ expression initiated in multipotent neural crest cells drives aggressive melanoma of the central nervous system. Pigment Cell Melanoma Res. 33, 96–111.

2. van der Kooij M.K., Speetjens F.M., van der Burg S.H., and Kapiteijn E. (2019). Uveal Versus Cutaneous Melanoma; Same Origin, Very Distinct Tumor Types. Cancers (Basel). 11,

3. Van Raamsdonk C.D., Bezrookove V., Green G., Bauer J., Gaugler L., O’Brien J.M., Simpson E.M., Barsh G.S., and Bastian B.C. (2009). Frequent somatic mutations of GNAQ in uveal melanoma and blue naevi. Nature. 457, 599–602.

4. Van Raamsdonk C.D., Griewank K.G., Crosby M.B., Garrido M.C., Vemula S., Wiesner T., Obenauf A.C., Wackernagel W., Green G., Bouvier N. et al. (2010). Mutations in GNA11 in Uveal Melanoma. N Engl J Med.

5. Carvajal R.D., Schwartz G.K., Tezel T., Marr B., Francis J.H., and Nathan P.D. (2017). Metastatic disease from uveal melanoma: treatment options and future prospects. Br J Ophthalmol. 101, 38–44.

6. Clarijs R., Schalkwijk L., Ruiter D.J., and de Waal R.M. (2001). Lack of lymphangiogenesis despite coexpression of VEGF-C and its receptor Flt-4 in uveal melanoma. Invest Ophthalmol Vis Sci. 42, 1422–1428.

7. Krantz B.A., Dave N., Komatsubara K.M., Marr B.P., and Carvajal R.D. (2017). Uveal melanoma: epidemiology, etiology, and treatment of primary disease. Clin Ophthalmol. 11, 279–289.

8. Clarijs R., Otte-Holler I., Ruiter D.J., and de Waal R.M. (2002). Presence of a fluid-conducting meshwork in xenografted cutaneous and primary human uveal melanoma. Invest Ophthalmol Vis Sci. 43, 912–918.

9. Mihic-Probst D., Ikenberg K., Tinguely M., Schraml P., Behnke S., Seifert B., Civenni G., Sommer L., Moch H., and Dummer R. (2012). Tumor cell plasticity and angiogenesis in human melanomas. PLoS One. 7, e33571.

10. Anand K., Roszik J., Gombos D., Upshaw J., Sarli V., Meas S., Lucci A., Hall C., and Patel S. (2019). Pilot Study of Circulating Tumor Cells in Early-Stage and Metastatic Uveal Melanoma. Cancers (Basel). 11,

11. Callejo S.A., Antecka E., Blanco P.L., Edelstein C., and Burnier M.N. (2006). Identification of circulating malignant cells and its correlation with prognostic factors and treatment in uveal melanoma. A prospective longitudinal study. Eye. 21, 752–759.

12. Keilholz U., Goldin-Lang P., Bechrakis N.E., Max N., Letsch A., Schmittel A., Scheibenbogen C., Heufelder K., Eggermont A., and Thiel E. (2004). Quantitative detection of circulating tumor cells in cutaneous and ocular melanoma and quality assessment by real-time reverse transcriptase-polymerase chain reaction. Clin Cancer Res. 10, 1605–1612.

13. Onken M.D., Mooren O.L., Mukherjee S., Shahan S.T., Li J., and Cooper J.A. (2014). Endothelial monolayers and transendothelial migration depend on mechanical properties of the substrate. Cytoskeleton (Hoboken). 71, 695–706.

14. Onken M.D., Li J., and Cooper J.A. (2014). Uveal Melanoma Cells Utilize a Novel Route for Transendothelial Migration. PLoS One. 9, e115472.

15. Harbour J.W., Onken M.D., Roberson E.D., Duan S., Cao L., Worley L.A., Council M.L., Matatall K.A., Helms C., and Bowcock A.M. (2010). Frequent mutation of BAP1 in metastasizing uveal melanomas. Science. 330, 1410–1413.

16. Yen M., Qi Z., Chen X., Cooper J.A., Mitra R.D., and Onken M.D. (2018). Transposase mapping identifies the genomic targets of BAP1 in uveal melanoma. BMC Med Genomics. 11, 97.

17. Schmidt S., and Debant A. (2014). Function and regulation of the Rho guanine nucleotide exchange factor Trio. Small GTPases. 5, e29769.

18. Feng X., Degese M.S., Iglesias-Bartolome R., Vaque J.P., Molinolo A.A., Rodrigues M., Zaidi M.R., Ksander B.R., Merlino G., Sodhi A. et al. (2014). Hippo-independent activation of YAP by the GNAQ uveal melanoma oncogene through a trio-regulated rho GTPase signaling circuitry. Cancer Cell. 25, 831–845.

19. Feng X., Arang N., Rigiracciolo D.C., Lee J.S., Yeerna H., Wang Z., Lubrano S., Kishore A., Pachter J.A., König G.M. et al. (2019). A Platform of Synthetic Lethal Gene Interaction Networks Reveals that the GNAQ Uveal Melanoma Oncogene Controls the Hippo Pathway through FAK. Cancer Cell. 35, 457–472.e5.

20. Nobes C.D., and Hall A. (1995). Rho, rac, and cdc42 GTPases regulate the assembly of multimolecular focal complexes associated with actin stress fibers, lamellipodia, and filopodia. Cell. 81, 53–62.

21. Parri M., and Chiarugi P. (2010). Rac and Rho GTPases in cancer cell motility control. Cell Commun Signal. 8, 23.

22. Nobes C.D., and Hall A. (1999). Rho GTPases control polarity, protrusion, and adhesion during cell movement. J Cell Biol. 144, 1235–1244.

23. Haga R.B., and Ridley A.J. (2016). Rho GTPases: Regulation and roles in cancer cell biology. Small GTPases. 7, 207–221.

24. Bokoch G.M. (2003). Biology of the p21-activated kinases. Annu Rev Biochem. 72, 743–781.

25. Zhao Z.S., and Manser E. (2012). PAK family kinases: Physiological roles and regulation. Cell Logist. 2, 59–68.

26. Guilluy C., Garcia-Mata R., and Burridge K. (2011). Rho protein crosstalk: another social network. Trends Cell Biol. 21, 718–726.

27. Acton S.E., Farrugia A.J., Astarita J.L., Mourão-Sá D., Jenkins R.P., Nye E., Hooper S., van Blijswijk J., Rogers N.C., Snelgrove K.J. et al. (2014). Dendritic cells control fibroblastic reticular network tension and lymph node expansion. Nature. 514, 498–502.

28. Sanz-Moreno V., Gadea G., Ahn J., Paterson H., Marra P., Pinner S., Sahai E., and Marshall C.J. (2008). Rac activation and inactivation control plasticity of tumor cell movement. Cell. 135, 510–523.

29. Chrzanowska-Wodnicka M., and Burridge K. (1996). Rho-stimulated contractility drives the formation of stress fibers and focal adhesions. J Cell Biol. 133, 1403–1415.

30. Byrne K.M., Monsefi N., Dawson J.C., Degasperi A., Bukowski-Wills J.C., Volinsky N., Dobrzyński M., Birtwistle M.R., Tsyganov M.A., Kiyatkin A. et al. (2016). Bistability in the Rac1, PAK, and RhoA Signaling Network Drives Actin Cytoskeleton Dynamics and Cell Motility Switches. Cell Syst. 2, 38–48.

31. Sander E.E., ten Klooster J.P., van Delft S., van der Kammen R.A., and Collard J.G. (1999). Rac downregulates Rho activity: reciprocal balance between both GTPases determines cellular morphology and migratory behavior. J Cell Biol. 147, 1009–1022.

32. Moore A.R., Ceraudo E., Sher J.J., Guan Y., Shoushtari A.N., Chang M.T., Zhang J.Q., Walczak E.G., Kazmi M.A., Taylor B.S. et al. (2016). Recurrent activating mutations of G-protein-coupled receptor CYSLTR2 in uveal melanoma. Nat Genet. 48, 675–680.

33. Onken M.D., Makepeace C.M., Kaltenbronn K.M., Kanai S.M., Todd T.D., Wang S., Broekelmann T.J., Rao P.K., Cooper J.A., and Blumer K.J. (2018). Targeting nucleotide exchange to inhibit constitutively active G protein α subunits in cancer cells. Sci Signal. 11,

34. Wyckoff J.B., Pinner S.E., Gschmeissner S., Condeelis J.S., and Sahai E. (2006). ROCK- and myosin-dependent matrix deformation enables protease-independent tumor-cell invasion in vivo. Curr Biol. 16, 1515–1523.

35. Nolen B.J., Tomasevic N., Russell A., Pierce D.W., Jia Z., McCormick C.D., Hartman J., Sakowicz R., and Pollard T.D. (2009). Characterization of two classes of small molecule inhibitors of Arp2/3 complex. Nature. 460, 1031–1034.

36. Matatall K.A., Agapova O.A., Onken M.D., Worley L.A., Bowcock A.M., and Harbour J.W. (2013). BAP1 deficiency causes loss of melanocytic cell identity in uveal melanoma. BMC Cancer. 13, 371.

37. Friedl P., and Wolf K. (2003). Proteolytic and non-proteolytic migration of tumour cells and leucocytes. Biochem Soc Symp. 277–285.

38. Sahai E., and Marshall C.J. (2003). Differing modes of tumour cell invasion have distinct requirements for Rho/ROCK signalling and extracellular proteolysis. Nat Cell Biol. 5, 711–719.

39. Sanz-Moreno V., Gaggioli C., Yeo M., Albrengues J., Wallberg F., Viros A., Hooper S., Mitter R., Féral C.C., Cook M. et al. (2011). ROCK and JAK1 signaling cooperate to control actomyosin contractility in tumor cells and stroma. Cancer Cell. 20, 229–245.

40. Rullo J., Becker H., Hyduk S.J., Wong J.C., Digby G., Arora P.D., Cano A.P., Hartwig J., McCulloch C.A., and Cybulsky M.I. (2012). Actin polymerization stabilizes α4β1 integrin anchors that mediate monocyte adhesion. J Cell Biol. 197, 115–129.

41. Collins C., and Nelson W.J. (2015). Running with neighbors: coordinating cell migration and cell-cell adhesion. Curr Opin Cell Biol. 36, 62–70.

42. Collins C., Denisin A.K., Pruitt B.L., and Nelson W.J. (2017). Changes in E-cadherin rigidity sensing regulate cell adhesion. Proc Natl Acad Sci U S A. 114, E5835–E5844.

43. Onken M.D., Worley L.A., Ehlers J.P., and Harbour J.W. (2004). Gene expression profiling in uveal melanoma reveals two molecular classes and predicts metastatic death. Cancer Res. 64, 7205–7209.

44. Humphries B., Wang Z., Li Y., Jhan J.R., Jiang Y., and Yang C. (2017). ARHGAP18 Downregulation by miR-200b Suppresses Metastasis of Triple-Negative Breast Cancer by Enhancing Activation of RhoA. Cancer Res. 77, 4051–4064.

45. Maeda M., Hasegawa H., Hyodo T., Ito S., Asano E., Yuang H., Funasaka K., Shimokata K., Hasegawa Y., Hamaguchi M. et al. (2011). ARHGAP18, a GTPase-activating protein for RhoA, controls cell shape, spreading, and motility. Mol Biol Cell. 22, 3840–3852.

46. Taniuchi K., Furihata M., Naganuma S., and Saibara T. (2018). ARHGEF4 predicts poor prognosis and promotes cell invasion by influencing ERK1/2 and GSK-3α/β signaling in pancreatic cancer. Int J Oncol. 53, 2224–2240.

