## Supplementary figures and images for "Uveal melanoma cells use ameboid and mesenchymal mechanisms of cell motility crossing the endothelium"

### Supplemental Figure S1

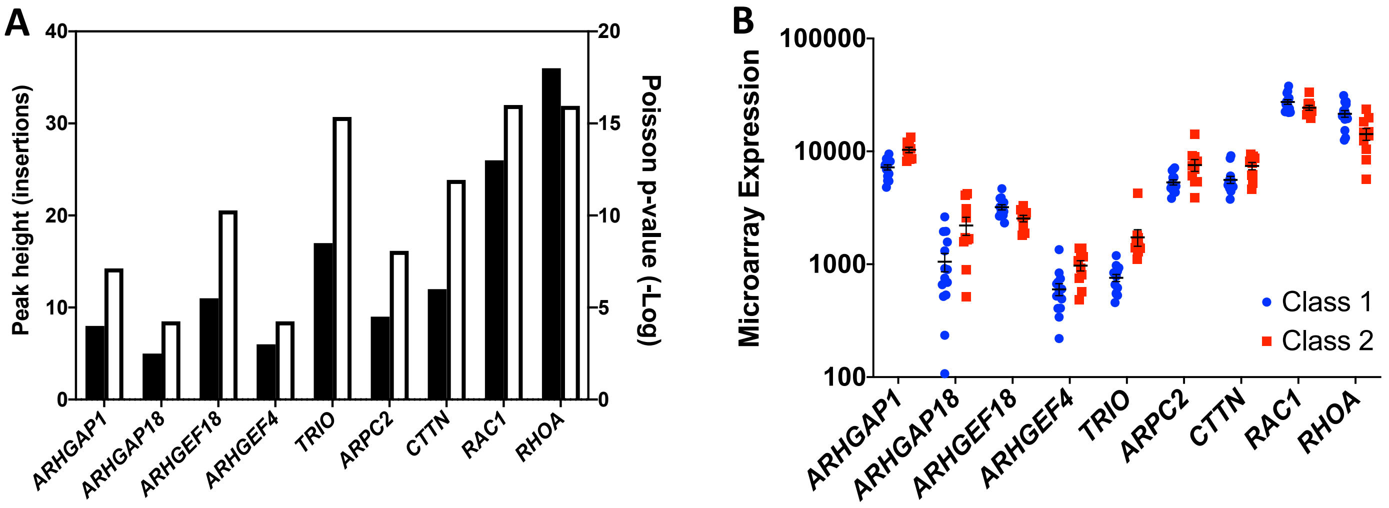

### Supplemental Figure S2

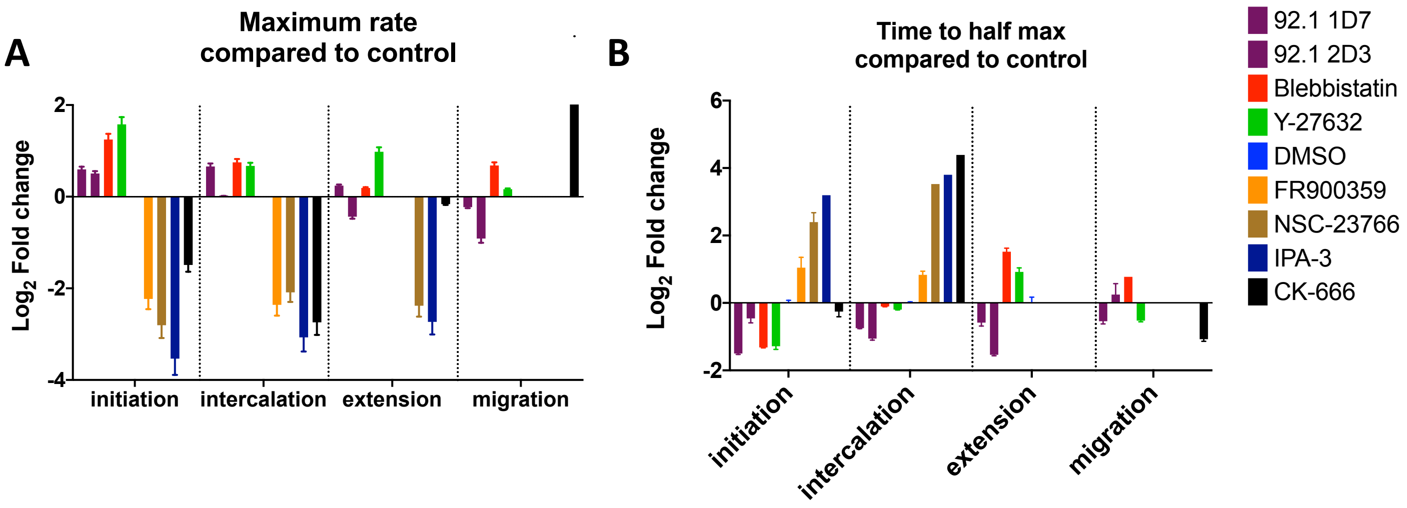
